# Diversified physiological sensory input connectivity questions the existence of distinct classes of spinal interneurons

**DOI:** 10.1101/2021.07.16.452238

**Authors:** Matthias Kohler, Fredrik Bengtsson, Philipp Stratmann, Florian Röhrbein, Alois Knoll, Alin Albu-Schäffer, Henrik Jörntell

## Abstract

The spinal cord is engaged in all forms of motor performance but its functions are far from understood. Because network connectivity defines function, we explored the connectivity for muscular, tendon and tactile sensory inputs among a wide population of spinal interneurons in the lower cervical segments. Using low noise intracellular whole cell recordings in the decerebrated, nonanesthetized cat *in vivo*, we could define mono-, di-, trisynaptic inputs as well as the weights of each input. Whereas each neuron had a highly specific input, and each indirect input could moreover be explained by inputs in other recorded neurons, we unexpectedly also found the input connectivity of the spinal interneuron population to form a continuum. Our data hence contrasts with the currently widespread notion of distinct classes of interneurons. We argue that this suggested diversified physiological connectivity, which likely requires a major component of circuitry learning, implies a more flexible functionality.

**Graphical Abstract:** 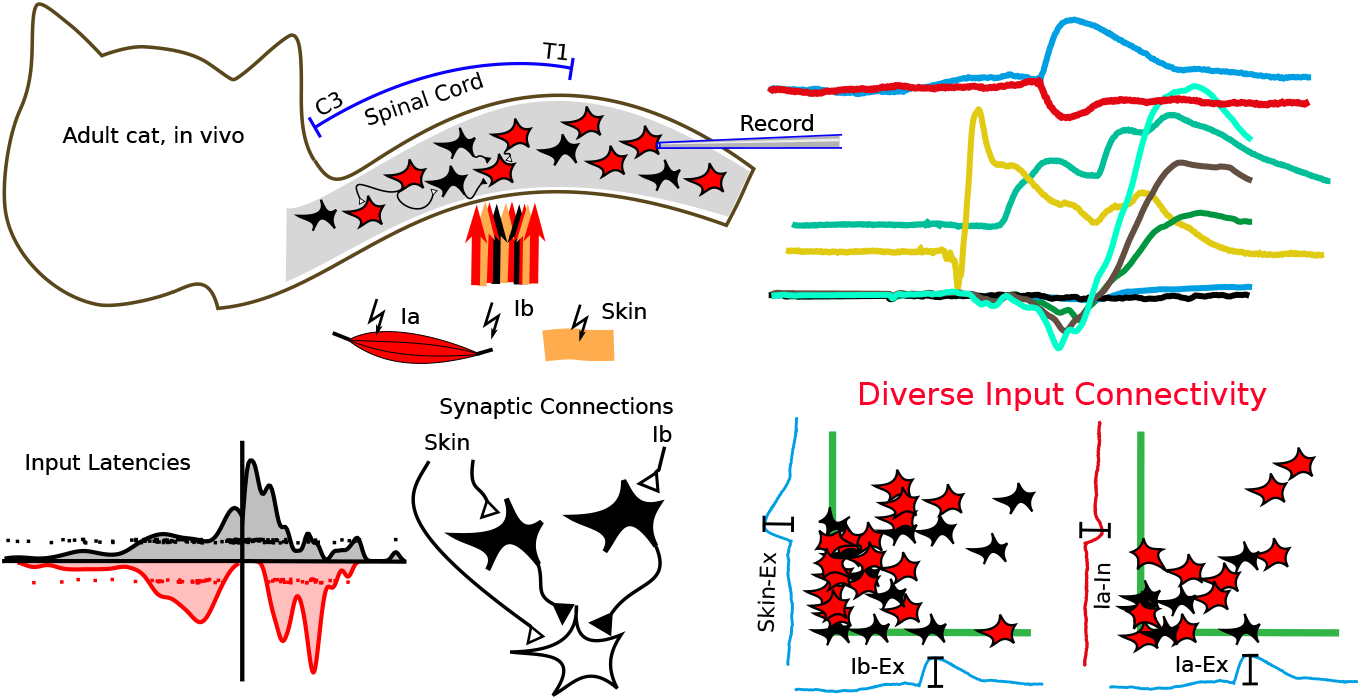

**Highlights:**

- *In vivo* whole cell, intracellular recording of spinal interneurons.
- Patterns of input from Ia, Ib and cutaneous afferents is highly diversified.
- Learning appears to be a defining factor of spinal interneuron connectivity.

## 3. Introduction

The spinal cord contains a potentially highly complex but poorly understood neuronal circuitry. Because of its position, it is pivotal to any aspect of brain action that involves movement (Crone et al., 2008; Bui et al., 2013; Azim et al., 2014) and its potential contributions to brain function can be considered underestimated (Loeb and Tsianos, 2015). In addition, because of the tight linkage between cognitive development and the development of movement (Diamond, 2000; Chiaravalloti et al., 2020), the functional principles of the spinal cord circuitry are potentially fundamental to many higher order functions by virtue of impacting how the brain perceives the world (Rongala et al., 2018).

The spinal cord has motoneurons, which are connected to the muscles, and interneurons, which are not. Major efforts have been directed to elucidate the organization of the spinal interneuronal network. One of the first indications of that there may be different types of spinal interneurons was the identification of the Renshaw cells, which receive input from motoneurons and inhibit motoneurons (Eccles et al., 1954). Later, the Ia inhibitory interneuron type was identified as a set of neurons receiving input from Ia afferents and inhibiting motoneurons of antagonistic muscles as well as other Ia inhibitory interneurons (Hultborn et al., 1971). Subsequent research focused on identifying additional types of spinal interneurons, by their input, by which connectivity pathway that the input arrived at the neuron, by their location and by their efferent synaptic action (Reviewed by Jankowska (1992) and McCrea (1992)). Though we are not aware of a publication that explicitly describes a classification scheme that assigns all spinal interneurons into disjoint classes based on their connectivity, this type of attempted neurophysiological classification approach had the aim of identifying all the specific interneuron types that could exist (McCrea, 1992). The implied goal of this major research endeavor was that eventually the resulting gigantic network puzzle of how all spinal interneurons are connected was going to be solvable. More recently, some authors have indicated that the identified classical types of spinal interneurons may not form distinct subsets, since for example some types clearly share some connectivity features with other types (Jankowska and Edgley, 2010; Hultborn, 2001).

Following in the footsteps of this major research endeavor of classical neurophysiology, spinal interneurons were found to be separable also on basis of their patterns of gene expression in early development (Jessell, 2000; Goulding, 2009). More than fourteen different main classes of spinal interneurons have been identified on the basis of differences in gene expression (Zholudeva et al., 2021). Recent studies suggested that the patterns of input connectivity, and thereby also the network formation, could be based on genetic preprogramming, where chemoattractants emitted by one interneuron type would attract synaptic inputs from a specific input source or specific interneuron types, suggesting that the neurophysiological and genetic classification schemes could be mapped to each other (Chen et al., 2006; Pecho-Vrieseling et al., 2009; Bikoff, 2019).

Here we investigated the input connectivity of spinal interneurons throughout the lower cervical spinal cord using intracellular recordings to define the synaptic linkage of their excitatory and inhibitory synaptic inputs from peripheral input sources in vivo. Our findings suggest an absence of distinct rules of input connectivity, and that every spinal interneuron in principle could be unique in this respect. We argue that in adult mammals the input connectivity, and therefore the function of the spinal cord circuitry, could to a large extent be defined by learning.

## 4. Results

We made 114 intracellular recordings of interneurons in spinal segments C6-T1 of the decerebrated cat (Figure 1 A). Neurons were recorded at depths between 1.31mm to 3.92 mm from the dorsal surface of the spinal cord, corresponding to anatomical laminae III-VII (Figure 1 B). Of these recordings, 68 were in the whole cell mode, where the recording quality allowed the measurement of the amplitude of individual synaptic responses. Thus we could analyze the synaptic weight. Responses were evoked by stimulation of the deep radial nerve and the skin. For the deep radial nerve, different stimulation intensities were explored, expressed as multiples of the stimulation threshold *T*, so that input from Ia and Ib afferents could be identified. The cutaneous afferents were stimulated to activate A-*β* fibers from the defined cutaneous receptive field of the neuron. The cells were recorded with mild hyperpolarizing currents of 10 pA to 30 pA to prevent spiking, thereby facilitating the analysis of the synaptic responses evoked from the peripheral sensory inputs. The general membrane physiology properties, from a subset of the neurons recorded in the whole cell mode, were previously reported (Spanne et al., 2014).

**Figure 1.**
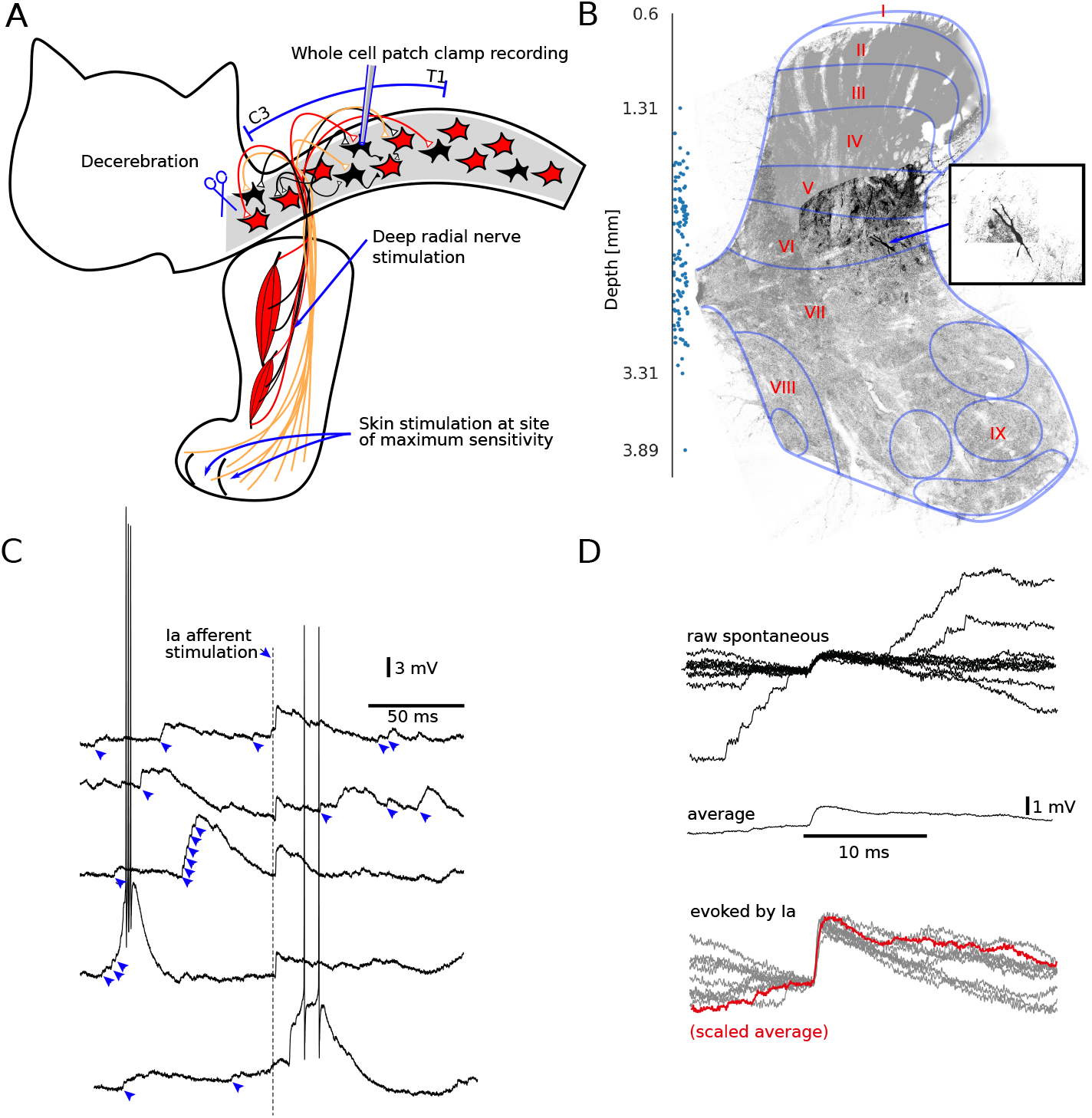
Experimental setup, location of recorded neurons and unitary EPSPs. **A** *In vivo* preparation, sites of stimulations and recordings. **B** Histological recovery of one recorded neuron with its location in the spinal cord grey matter and a detail image of the recorded neuron with a visible soma and proximal dendrites. The image was acquired by a confocal microscope from a 50 μm thick section from the spinal segment C8. Rexed laminae I-IX are tentatively indicated (Rexed, 1952). Additional plot of the depths of each recorded neuron with a random offset in the medial-lateral axis. **C** Five raw traces of recordings from a spinal interneuron (approximate resting *V_m_* = −60 mV) with near threshold DR stimulation (’Ia afferent stimulation’ at dashed vertical line). Blue arrowheads indicate prominent spontaneous EPSP-like responses. The raw traces also illustrate examples of spontaneous spiking, which was reduced by mild hyperpolarizing currents throughout recordings to facilitate the analysis of the evoked synaptic responses. **D** Superimposed spontaneous EPSPs, their average and a comparison between the latter and superimposed EPSP responses evoked by Ia afferent stimulation (DR stimulation at 1.05 times threshold).

### 4.1. Matching the shape of the recorded potentials to synaptic activations

Decerebration allowed the recordings to be made *in vivo* without anesthesia. This kind of animal model is a system with spontaneous activity in both the spinal neurons and the sensory afferents. The intracellularly recorded activity was characterized by a high number of spontaneous events (Figure 1 C). These events occasionally summed to sufficient depolarization to trigger discharge of an action potential in the recorded neuron. The spontaneous events were typically found to have a shape that was congruent with other spontaneous events, and they all displayed a voltage-time shape similar to unitary excitatory post synaptic potentials (EPSPs) recorded in other parts of the central nervous system (Jörntell and Ekerot, 2006; Bengtsson and Jörntell, 2009; Bengtsson et al., 2013). When multiple such events were averaged, it was more clear that they had the typical shape of an EPSP (Figure 1 D). Unitary responses evoked using electrical stimulation of peripheral afferents at a sufficiently low intensity (near activation threshold) had the same shape. These findings show that the spontaneous events with this shape were EPSPs, which could be triggered by spontaneously active external afferents or other spinal interneurons by electrical activation of peripheral sensory afferents.

### 4.2. Evoked compound PSP responses and mapping of the input connectivity

For each recorded cell we could identify unitary inhibitory postsynaptic potentials (IPSPs) and EPSPs (Figure 1 D). Averaging at least one-hundred such PSPs, for each recorded neuron, was used to create template EPSPs and IPSP, respectively (Figure 2 A). We observed that the time constants of template PSPs could differ somewhat between different neurons, but the differences were not dramatic. Similarly, within each neuron, different spontaneous EPSPs could potentially differ somewhat between different EPSP events, but the template was designed to correspond to the average of these events. Using scaling of the template PSPs, we could reconstruct the raw PSP responses evoked by the afferent stimulation. The scaling of the template PSPs was done to capture the shape of the evoked PSPs also when they were due to the activation of multiple synapses at the same time (Bengtsson et al., 2013). Pure monosynaptic EPSPs were readily identified by their distinct overlap with the template PSP (Figure 1 D).

**Figure 2.**
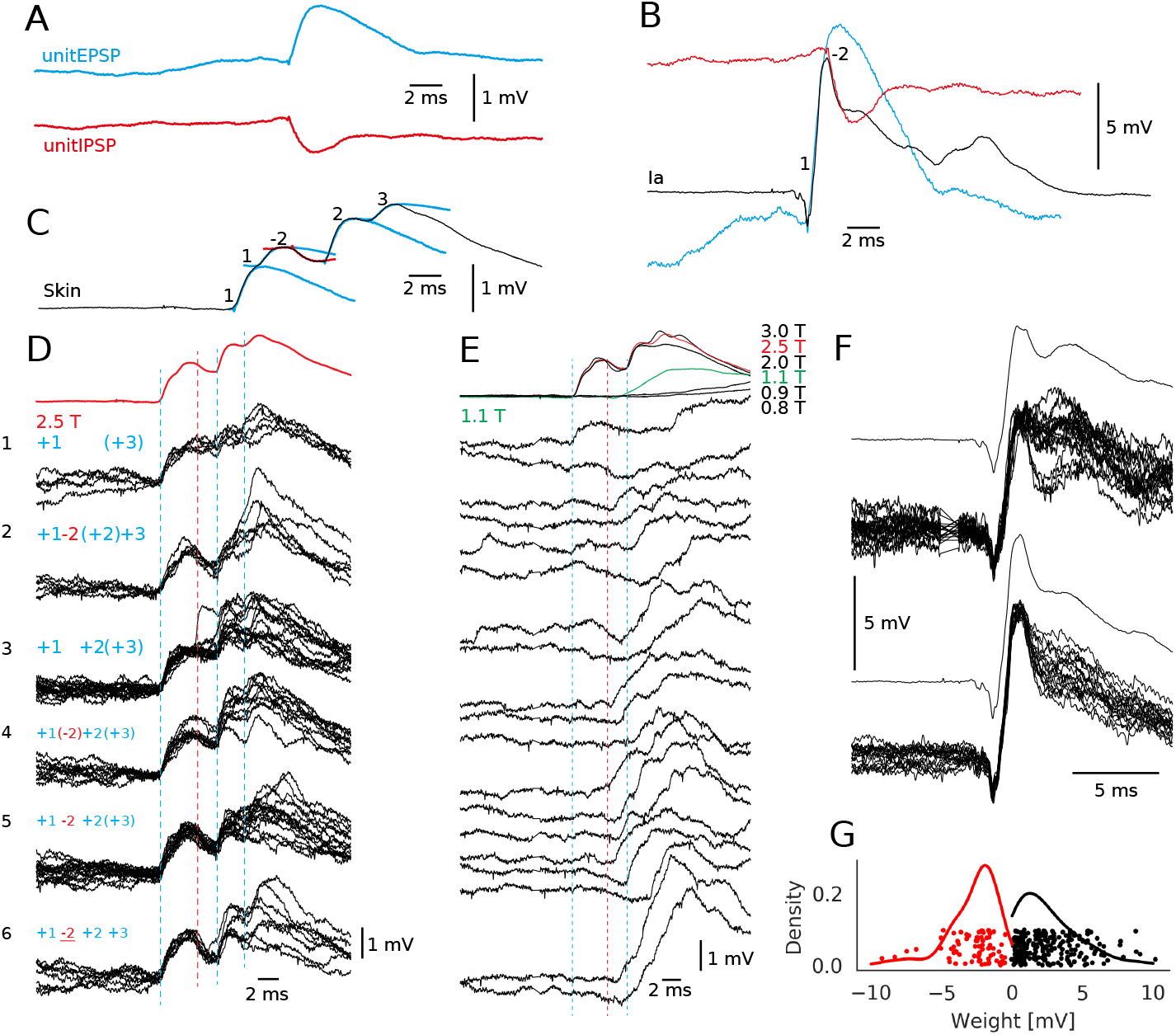
Definition of input connectivity based on template reconstruction of sensory-evoked compound synaptic responses. **A** Excitatory post synaptic potentials (EPSPs) were averaged to obtain a unit EPSP template. The same was done for inhibitory PSPs (IPSPs). These templates were used to decompose compound evoked synaptic responses. **B** An average PSP response evoked by stimulation of the deep radial nerve (black) that could be decomposed into a (monosynaptic, ”1”) EPSP and a disyanptic IPSP (”−2”) based on amplitude scaling of the PSP templates. **C** Decomposition of an averaged compound synaptic response evoked by skin stimulation. In this sample cell, the response was decomposed into a mono-, di- and trisynaptic EPSP and a disynaptic IPSP. The numbers indicate whether the input was mono-, di-, or trisynaptic, the sign indicates whether it was excitatory or inhibitory **D** Average (red trace) and raw responses evoked by skin stimulation at 2.5*T*. Raw traces are grouped by their relative similarity. Brackets around numbers indicate that the specific input was weak, whereas underscored input indicates that the input was strong. **E** Average and raw responses evoked by skin stimulation at 1.1*T*. **F** Average and raw responses evoked by DR stimulation. Note that the IPSP partially disappears at the lower DR stimulation intensity. **G** Kernel density estimations of the synaptic weight distributions of excitatory (red) and inhibitory (black) PSPs across all neurons recorded. Additional plot of individual weights with a random offset in the density axis, and a random offset from 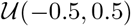 in the weight axis.

Compound raw responses, resulting from the activation of partly overlapping EPSPs and IPSPs, were analyzed by a template fitting process. The template fitting process consisted of scaling the template PSP to the observed response, which thereby yielded a response amplitude and a response latency time for that evoked event. In Figure 2 B and C, example results of the fitting process are illustrated for responses evoked by skin input and muscle afferent input, respectively. Since the responses of this neuron to the two inputs were relatively consistent, we could in this illustration apply the template fitting even to the average responses. The analysis was however applied to individual raw traces, using the same principles as illustrated in Figure 2 B,C. In Figure 2 D-F, we show how this method was applied to the single raw traces. In Figure 2 D, the exact same stimulus intensity (2.5*T*, average response on top, in red, same as in Figure 2 C) evoked six main groups of responses. The earliest part was the monosynaptic response component (+1, first blue dashed vertical line). The monosynaptic response actually consisted of two separable components, likely because it contained afferents of two separate mean conduction velocities (Bengtsson et al., 2013). The monosynaptic response was essentially invariant across the six groups, i.e. it was evoked at the exact same response latency time with a response amplitude with low variance.

In comparison, di- and trisynaptic responses were more variable with respect to both amplitude and latency (Figure 2 D and also Figure 4 A,B). This is to be expected for non-monosynaptic inputs as the transmission is dependent on the (time-varying) excitability of the intercalated spinal interneurons at the time of sensor activation. Moreover, the intercalated neurons will also lead to the response onset latency times being substantially longer than for the monosynaptic response, as measured from the time of arrival of the afferent input into the spinal cord white matter. This can for example be seen for the disynaptic IPSP (−2, average onset latency time at the red dashed vertical line) which was essentially absent from the responses in groups 1 and 3, and much more prominent in group 6. Later response components (+2 and +3) were also variable in their occurrence, amplitude and response latency times. Figure 2 E illustrates the raw responses to a stimulation intensity of just above the threshold (1.1*T*). In this case, the monosynaptic EPSP and the disynaptic IPSP essentially did not occur, but the more indirect excitatory synaptic responses remained. In this case, the wildly variable nature of these indirect responses were more clear. Figure 2 F illustrates the raw responses evoked from the deep radial nerve (DR) at two different stimulation intensities. At the lower intensity (1.1*T*), the disynaptic IPSP was highly variable, whereas at the higher intensity (1.2*T*), it was more consistently present. In contrast, the monosynaptic EPSP remained constant across both intensities, even though the higher intensity evoked a compound EPSP with a somewhat larger amplitude. For further analysis, synaptic weights were measured for each input connection as the average response amplitude. Figure 2 G shows the distribution of these weights across all neurons where the recording quality allowed the amplitude to be measured (236 excitatory and 74 inhibitory inputs in 68 neurons). The weight distribution is skewed, which is a commonly observed phenomenon among synaptic weights that have been subject to learning (Barbour et al., 2007).

We used stimulation of the deep radial nerve, which contains no skin afferents, to stimulate the Ia and the Ib afferents of the supplied muscles, as previously described(Quevedo et al., 2000). Ia and Ib responses could be separated, even though both inputs were activated through the same nerve. The identification of whether a recorded neuron was receiving synaptic inputs from Ia or Ib afferents, or both which did occur in some cases, was based on several criteria similar to those previously described in the literature. With nerves from the cat hind limb there is often a difference in threshold as well as conduction velocity between Ia and Ib fibres (Bradley and Eccles, 1953; Eccles et al., 1957a,b; Laporte and Bessou, 1957; McCrea et al., 1995). Similar to these previous studies, we used field potential recordings in the extracellular tissue outside the recorded neuron to identify the threshold of the Ia volley, where the Ia volley corresponds to the field potential generated by the action potentials of a population of Ia afferents (Figure 3 A). Monosynaptic inputs could be defined on the basis of their very short delay relative to the local field potential (LFP), corresponding to the activation of the synapses on the spinal neurons, that was initiated about 0.5 ms after the onset of the Ia afferent volley. For the DR stimulation, a neuron activated at 1.0*T* – 1.1*T* was considered to be activated by Ia afferents (Figure 3 B), whereas a neuron that was activated distinctly only above 1.2*T*, typically well above this value, was considered to be activated by Ib afferents (Figure 3 C). In addition to the threshold criterion, for neurons activated monosynaptically, the responses latency times depended on whether they were activated by Ia or Ib afferents, as defined by the threshold criterion (Figure 3 C). These criteria could also be used to identify neurons which had monosynaptic input from both Ia and Ib (Figure 3 D). This convergent input is in agreement with a previous report based on more indirect observations of such convergence (Jankowska and McCrea, 1983).

**Figure 3.**
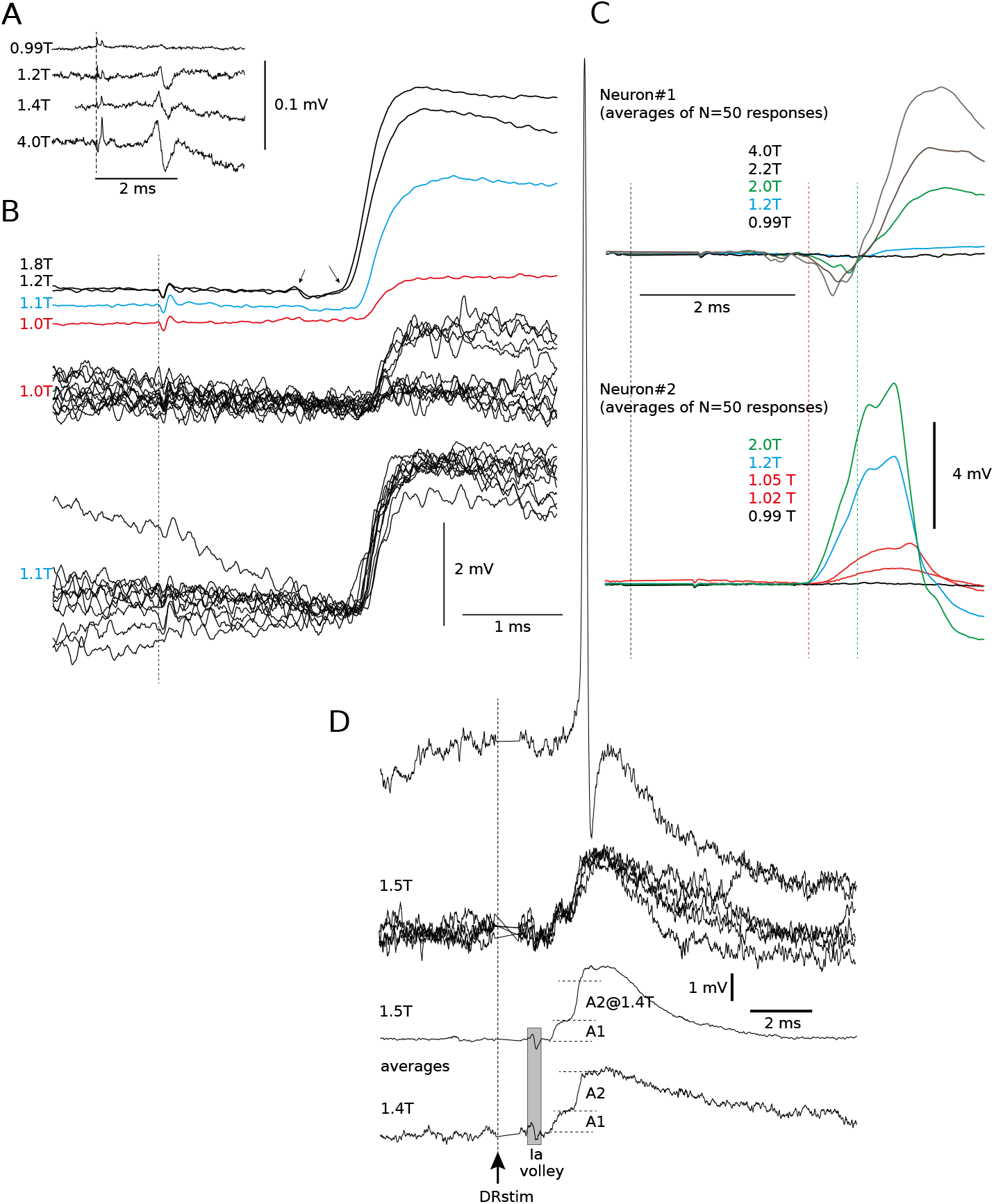
Separation of responses from Ia and Ib afferents activated from the deep radial (DR) nerve. Black vertical dashed lines indicate the onset of stimulation in all panels of this figure. **A** Definition of threshold intensity *T* for activating the Ia afferent nerve volley based on the local field potentials (LFP) recorded within the spinal grey matter, outside the recorded neurons. Note that the response latency shortens slightly with increased stimulation intensity. **B** From the same experiment after establishing an intracellular recording, EPSPs evoked at and above *T*. Top, averages of the responses evoked at four different intensities. Arrows indicate the Ia afferent volley and the onset of the EPSP response. Below, superimposed raw responses evoked at 1.0*T* and 1.1*T*. **C** From another experiment, average recordings of evoked EPSPs from two different neurons, recorded adjacently in the same electrode track. Neuron#1 responded only at intensities > 1.2*T*, and the EPSPs had a distinctly longer response latency time than the preceding Ia synaptic LFP (starting at the red dashed line). Bottom set of recordings, Neuron#2 responded at all intensities above 0.99*T*. Green dashed line indicates response latency time for the evoked EPSP in Neuron#1, red dashed line indicates response latency time for the evoked EPSP in Neuron#2. For Neuron#2, note that the stimulation at higher intensities sometimes evoked a spike, which created the ripple in the later part of those responses. Note that there was no clearly visible Ia volley in these two recordings. **D** Recording from a neuron with monosynaptic input from both Ia and Ib afferents. Top, raw traces evoked at 1.5*T*, one of which also contained an evoked spike when the clamping current was released for one of the simulation repetitions. Average of responses at 1.4*T* and 1.5*T* show two summed EPSPs with amplitudes A1 and A2. The amplitude of A1 was the same at 1.4*T* and 1.5*T* but the A2 amplitude was larger at 1.5*T*, which alongside the differences in latency times of the two inputs indicate that response A1 was a Ia EPSP and A2 was a Ib EPSP.

We also checked whether repeated activations of afferent input led to changes in the response amplitudes of the evoked PSPs. Over monosynaptic evoked PSPs that appeared at least twenty times at intervals of either 1 s or 334 ms, the average coefficient of variation of the response amplitude was 0.122 (*SD* = 0.041). Also, we could not find any significant evidence of depression or potentiation. A linear fit to the response amplitudes has an average increase of −0.001 mV per stimulation repetition (*SD* = 0.01250). We next analyzed the response latency times for each identified input, across all neurons. Figure 4 A-B shows that the responses with longer response latency times also tended to have a substantially higher variance in their response latency times (note the logarithmic scale in Figure 4 A-B). This is an indication of a response being mediated more indirectly, over multiple synapses. When analyzing the distribution of the response latency times of the PSPs evoked by the different types of afferents they tended to form clusters. For PSPs that were close to the cluster boundaries, their classifications were additionally confirmed by their stimulation threshold (Figure 3). This formed the basis for determining the input connectivity across all neurons recorded, whether the input connectivity was mono, di or tri-synaptic in addition to whether it was excitatory or inhibitory.

**Figure 4.**
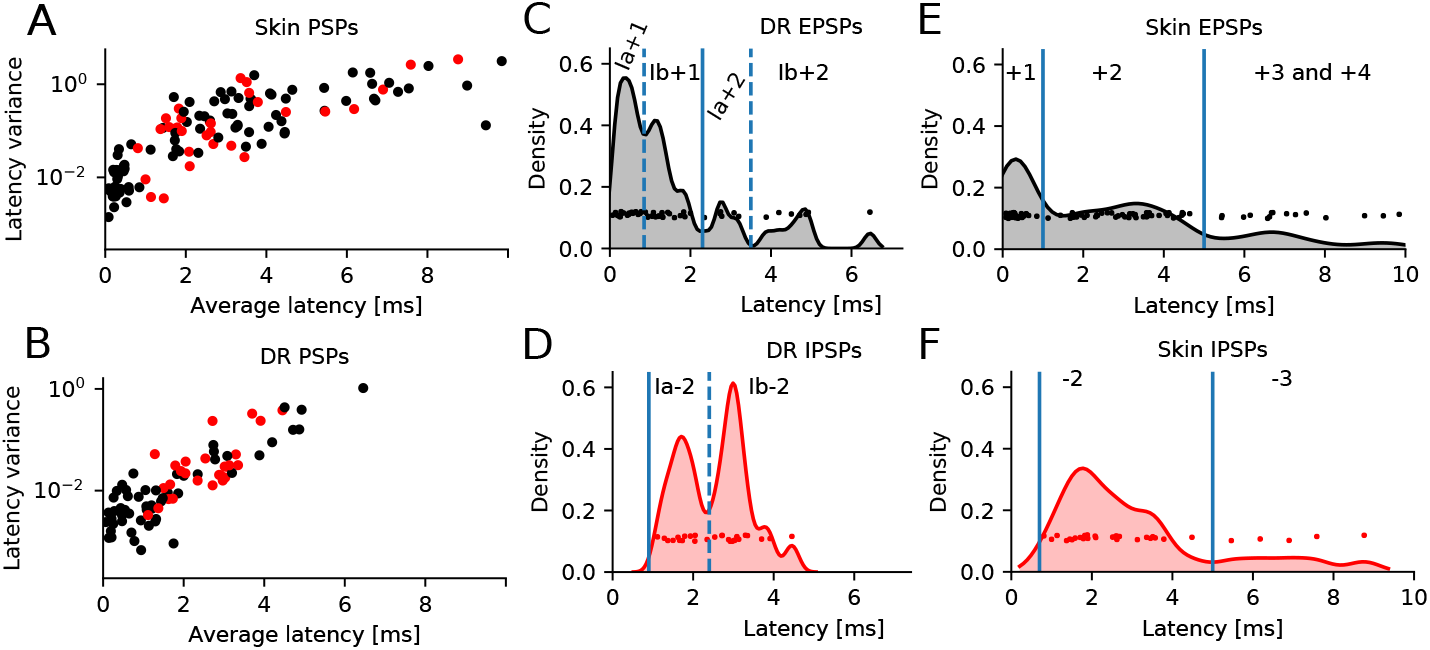
Definition of input connectivity based on response latency times. **A-B** The average response latency time plotted against variance of the response latency time for EPSP (red) and IPSP (black) responses. **C–F** The distribution of the average response latency times of the evoked PSPs for each recorded response component in each neuron (the data points are given a small random offset along the density axis for better visualization of the density). A kernel density estimation of the density of responses at each given response latency time is overlaid on the data points (0.1ms resolution in **C** and 0.2 ms resolution in **D-F**). Latency times for responses evoked by the DR and the skin stimulation were normalized by subtracting the latency time of the respective LFP. Separation of PSPs evoked by Ia and Ib afferents, respectively, is indicated by dashed blue lines.

### 4.3. Clusterability of neurons by input connectivity

We next proceeded by making a systematic analysis of the input connectivity of all recorded neurons. The purpose was to explore if the input connectivity across the neuron population formed clusters or classes. First we inspected plots of pairwise weighted input combinations. These input combinations were picked from all recorded inputs that were either monosynaptic excitation or disynaptic inhibition. No classification by input connectivity was immediately obvious (Figure 5 A), because none of the emerging distributions formed any clusters. Instead, they rather resembled a continuous single modal distribution. Hence, these findings did not indicate any specific rules of input connectivity.

**Figure 5.**
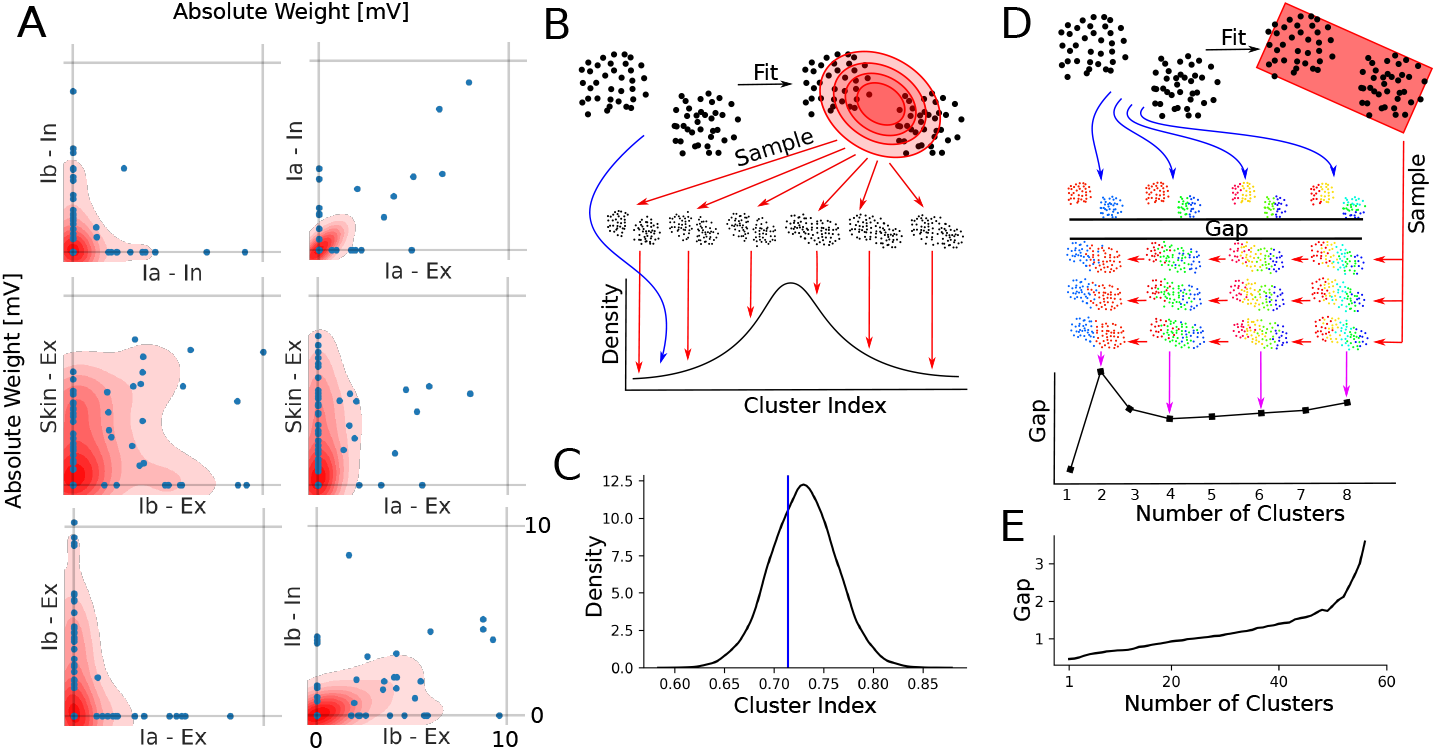
Absence of correlation patterns and clusterability of neurons by input combinations. **A** Kernel density estimations of absolute synaptic weights from distinct combinations of monosynaptic excitatory and disynaptic inhibitory inputs. (In: Inhibitory, Ex: Excitatory) No subdivision between the neurons is visible, though for some combinations (Ia-In, Ia-Ex) there appears to be some correlation. Additional plot of nonzero input weights at a random offset from 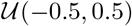. **B** Illustration of analysis method and the output of the SigClust test expected for a hypothetical dataset with clusters. The clustering index of the original dataset (blue arrow) is compared to that of random samples from a Gaussian distribution (red arrows) fitted to the dataset (black arrow). **C** The SigClust results for our set of recorded neurons. The clustering index for the dataset of recorded neurons (blue line) compared to the fitted Gaussian distribution (black). **D** Illustration of the GapStat analysis method and expected output for a hypothetical dataset with two clusters. The GapStat method first fits a uniform distribution to the data. The samples are drawn from that distribution (blue arrows) and for each possible number of clusters the gap to the actual dataset (red arrows) is computed. The number of clusters at which the gap to the actual dataset is maximized is the estimated number of clusters for the dataset. **E** The GapStat results for our set of recorded neurons. The plot illustrates that the gap just continues to grow with the number of tested clusters. Hence this method does not find our data clusterable.

In order to determine in a more standardized fashion if the spinal interneurons could be clustered on basis of their input connectivity, we next utilized the SigClust and GapStatistics tests on the part of the dataset where also the synaptic weights for each input could be defined (*n* = 68 neurons).

The SigClust test (Figure 5 B) can be used to test if a dataset can be split into at least two clusters by comparing the actual data to a number of random samples from a Gaussian distribution fitted to the dataset. If the original dataset has a significantly smaller clustering index (blue arrow in Figure 5) than random samples of the Gaussian fit (red arrows in Figure 5 B), then the SigClust test returns a probability that the dataset is containing clusters. Figure 5 C shows the result of this test on our dataset. The cluster index, which measures how well the dataset can be split into two clusters reported a value of 0.71. The average cluster index of the distribution of random sample datasets was 0.729 with a standard deviation of 0.033. Hence the hypothesis that the dataset was not clusterable could not be rejected (*p* = 0.31).

The GapStatistics test (Figure 5 D) is another method that can be used to test if a dataset can be clustered and in addition it estimates an appropriate number of clusters for the dataset. The appropriate number of clusters is estimated as the point where the gap measure between the dataset and random samples from a fitted uniform distribution is maximized (Figure 5 D). However for our dataset of recorded neurons the gap continued to increase with increasing number of clusters (Figure 5 E). When the gap just continues to grow with the number of datapoints added, as in this case, it indicates that the data does not contain any clusters. Hence, all of these three analyses indicated that our population of recorded interneurons did not contain any specific input connectivity classes.

### 4.4. Spinal network structure

Given that our data provided input connectivity data also for indirect, non-monosynaptic inputs, it implied that some of the recorded neurons could be responsible for mediating the indirect input to other recorded neurons. Hence, we also tested if the input connectivity observed across the full population of neurons (*n* = 114) could be explained by the pool of recorded input connectivity data. We first tested if the recorded input connectivity could be explained by a pure feed-forward network.

We did this by reconstructing a hypothetical feed forward network, for all recorded input combinations observed up to four sequential synapses away from the monosynaptic sensory input source. This analysis built on the principle shown in Figure 6 for three exemplary neurons, i.e. for any neuron receiving non-monosynaptic input, there must be predecessor neuron(s), which mediate(s) that input.

**Figure 6.**
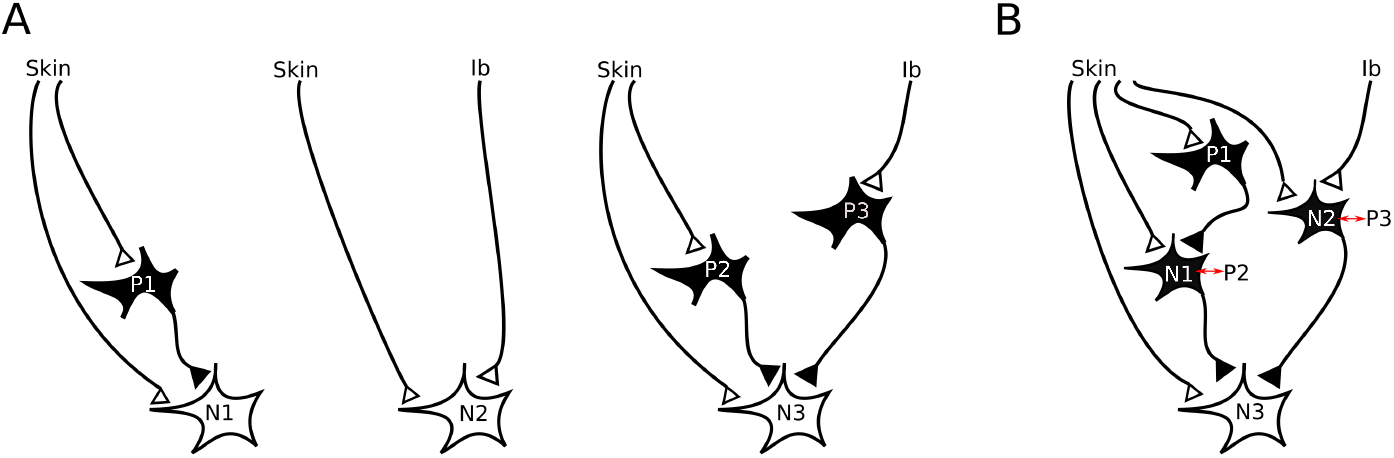
Matches between the input connectivity of individual target neurons and predecessor neurons explain all inputs observed. **(A)** Input connectivity of three recorded neurons *N*1, *N*2 and *N*3. **(B)** The input connectivity of neuron *N*3 can be explained by the input connectivity of neurons *N*1 and *N*2.

Figure 6 A shows the input connectivity for three of our recorded neurons *N*1, *N*2 and *N*3. Neuron *N*1 received excitatory monosynaptic input from Skin and disynaptic inhibitory input from Skin. For the inhibitory input an inhibitory interneuron (Neuron *P*1) is necessary. Neuron *N*2 received monosynaptic excitatory input from Skin and Ib. Neuron *N*3 received the same input as *N*1 plus disynaptic inhibition from Ib. The inhibitory input to *N*3 could then have been provided by the two inhibitory interneurons *P*2 and *P*3. The input connectivity of Neuron *N*3 can be explained by the input connectivity of neurons *N*1 and *N*2. If the neuron *N*1 is assumed to be an inhibitory interneuron, then it could correspond to neuron *P*2. If the neuron *N*2 is assumed to be an inhibitory interneuron, then it could correspond to neuron *P*3. (Figure 6 B).

Hence, the network connectivity pattern observed in these three recorded interneurons could be readily explained by our recording data. Note that the presence of *P*1 would not be possible to detect through the recording of *N*3, as disinhibitory effects were not detectable, but its presence could be detected by the *N*1 recording. For all but one of our recorded neurons, the input connectivity to each neuron could in each case be explained by a recorded predecessor neuron, where a valid predecessor contained the necessary input and in addition did not mediate input that was not observed in the recorded neuron. Therefore, all recorded patterns of input connectivity could be explained by monosynaptic inputs plus the inputs observed in other neurons in our data. Most of these connections were likely to be of a feed-forward type, although, due to the lack of input classes, there were no indications of any specific layers within the spinal cord circuitry.

Using a similar approach, we found that the number of potential disynaptic excitatory loop connections in our recorded connectivity data was significantly reduced (*p* < 0.05) compared to chance (Figure 7).

**Figure 7.**
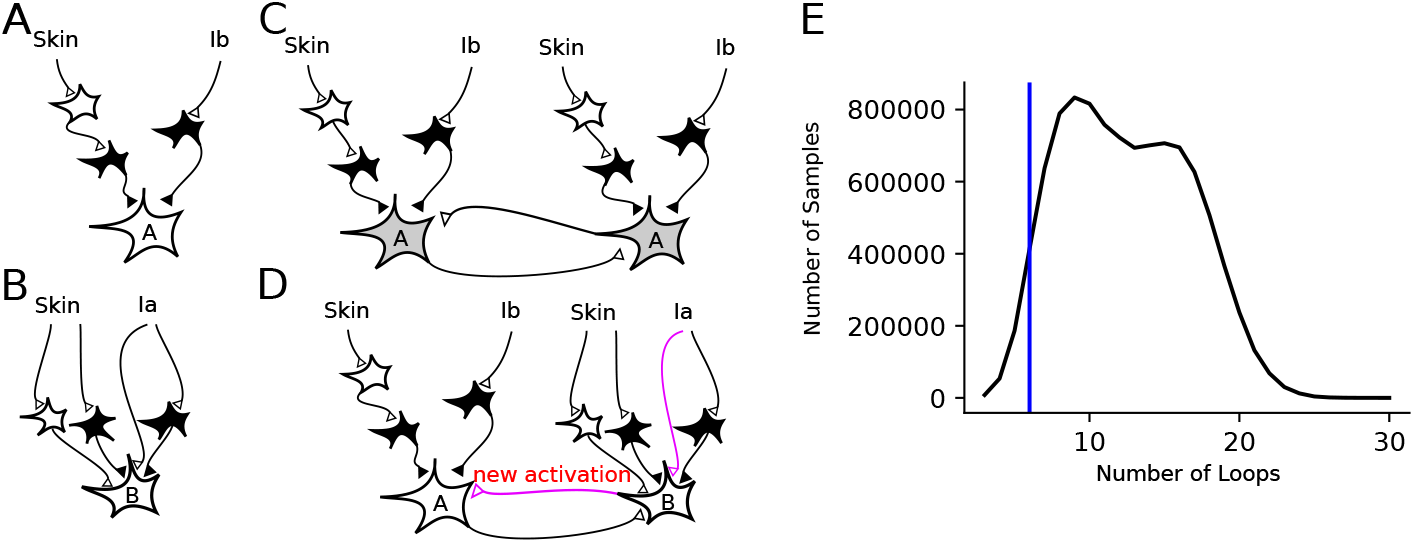
Disynaptic excitatory loops were suppressed compared to chance. **A,B** Two example input combinations. **C** Two neurons with input combination A could potentially be connected to each other, as our recording data could not detect inhibited synaptic excitation. **D** Neurons with input combinations A and B could not be connected to each other as the added disynaptic Ia excitation (as well as added trisynaptic skin excitation) would have been detected in our recording data. Hence, we can use our recording data to estimate the number of potential reciprocal excitatory loops. **E** Comparison of the observed number of potential excitatory loops in the recorded data (blue line) compared to the swap randomized datasets. Reciprocal excitation was significantly reduced (*p* < 0.05) compared to chance, i.e. the swap randomized dataset of connectivity.

### 4.5. Associations between neuronal inputs

Since correlations and anticorrelations in the input connectivity were visible in the analysis above (for instance a possible linear relation between Ia-Ex and Ia-In in Figure 5A), we searched for any possible correlation or anticorrelation in the inputs systematically, using an association rule analysis.

In our data, a spinal interneuron could receive input from the Ia, the Ib and the skin afferents as IPSPs and/or as EPSPs. Note that this part of the analysis ignored whether the input was mono-, di-, tri- or quadsynaptic. Since only the binary absence or presence of input was considered, and not the weights of the inputs, the full dataset of 114 neurons was used.

We tested if any of the resulting thirty possible input combinations (associations) were significantly enhanced or reduced compared to a random distribution (Figure 8). The random distribution was generated by swap randomizing the input combinations of our dataset (Figure 8 A). The position of the recorded data in relation to the distribution of this randomized data was used to indicate if the presence of each specific connectivity pattern was enhanced or reduced compared to chance connectivity.

**Figure 8.**
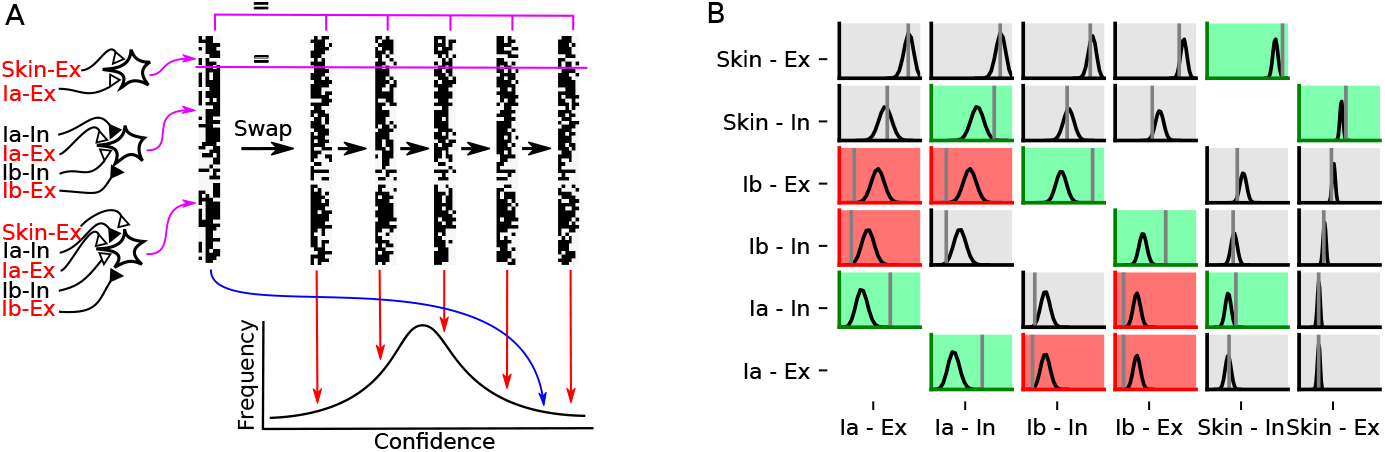
Estimation of the observed input combinations compared to chance. **A** The input connectivity of the recorded neurons is represented as a matrix of binary elements (left column) where each line represents the input to one recorded neuron. In this analysis, we only considered input from the three main sensory sources and whether the input was excitatory or inhibitory. Swap randomization was used to construct random matrices with equal number of connections as in the observed data for each respective neuron. The scrambled data is used to build a distribution of random connectivities (red arrows) against which the confidence of the recorded connectivity patterns can be compared (blue arrow). **B** For each possible combination of inputs, the confidence of the specific input combination of the data versus the distribution of confidence of the swap randomized data is shown. Associations that are significantly enhanced are marked green, significantly reduced associations are marked red. Input combinations that were not significantly enhanced or reduced are shown in grey. The individual significance tests are corrected for multiple comparisons such that the total significance level is at 0.05.

The threshold of significance for each test was adjusted for multiple comparisons so that the total significance of all tests was *p* = 0.05. For both the Ia afferents, the Ib afferents and the skin input, the associations between inhibition and excitation were significantly enhanced (Figure 8 B, green panels). The association between inhibitory input from the skin and Ia afferents were significantly enhanced (green panel). Some combinations of Ia and Ib afferent input were significantly reduced (Figure 8 B, red panels). None of the other input combinations were found to be reduced or enhanced at a significant level (grey panels). Hence, the association rule analysis suggested that overall there were only a few constraints on the probability of specific input combinations among our set of recorded spinal interneurons.

## 5 Discussion

We made intracellular recordings from a variety of spinal interneurons in the lower cervical segments *in vivo*. We recorded their synaptic input connectivity from a variety of input sources and defined the synaptic linkages by which the inputs were provided. The inputs to each individual spinal interneuron were well organized, with large to very large compound synaptic responses for the input that was present, but absent or very weak input for all other inputs. However across the population of spinal interneurons the synaptic input patterns formed a great diversity. This provided no support for specific sets of spinal interneuron input classes. Instead, the patterns of input connectivity to the spinal interneurons was better described as a continuum. The only exceptions were a reduced association between Ia and Ib excitation and an enhanced association between excitation and inhibition for inputs from the same source (Ia, Ib and skin). As discussed below, a spinal cord with a continuum of input combinations, rather than a limited set of fixed entities, indicates that much of the input connectivity is most likely shaped by learning. This carries the advantage of making the spinal cord circuitry better geared to cope with complex motor tasks and flexibility in motor control.

### 5.1. Methodological considerations

Since we kept the skin almost entirely intact, except for a small incision around the deep radial (DR) nerve, we could manually identify the skin area of maximum sensitivity for each recorded interneuron. It is known that each spinal interneuron can receive different types of synaptic inputs from different skin areas, i.e. excitation from one skin area and inhibition from another (Weng and Schouenborg, 1996). Therefore, we know that we have underestimated the complexity of the skin input connectivity to our neurons. Moreover, because of our requirement to keep the skin as intact as possible, an experimental shortcoming was that we could only use the DR nerve for stimulation of muscle afferents. The DR nerve supplies a high number of muscles in the forearm and paw, but not all. Hence, there were several muscles for which we could not test the presence of muscle afferent input. Also, we could not be certain that the inputs provided from Ia and Ib afferents, and excitatory versus inhibitory synaptic inputs, were from the same muscle or not. The interactions between antagonists and agonists could also not be studied. The skin inputs were limited to tactile receptors, which provide input over A-*β* fibers. These include rapidly and slowly adapting subtypes with various end organs. Hence, we again underestimated the diversity of skin inputs. Therefore, even though our data shows a highly complex connectivity, it is clear that our data represents an underestimate of that complexity. We only focused on the input connectivity. If we in addition had been able to take the output connectivity into account, i.e. what other neurons in the spinal cord and in supraspinal structures they were connected to (Geborek et al., 2013; Jörntell, 2017), then our population of recorded neurons would appear at least as diverse. Had we additionally taken into account factors, which are not directly related to motor control, such as the expression of cell adhesion molecules, then our population of neurons would also appear at least as diverse, if not more than described here.

Our *in vivo* intracellular recordings provided many advantages for the study of the physiological connectivity (anatomical connectivity combined with the weights of the synaptic inputs) of the fully developed mammalian spinal cord. First, decerebration rather than anesthesia provided the advantage that anesthesia-induced suppression of neural and synaptic activity was omitted and hence also low strength synaptic responses with multiple synaptic relays could be reliably recorded. The low recording noise of the whole cell patch clamp technique also provided substantial advantages regarding the resolution of the recordings. For example, even unitary synaptic potentials could be recorded, compared to the classical neurophysiological recordings where such weaker inputs would have been at substantially higher risk of being missed. The combination of these two methodological advantages could be a main explanation for why we could see a continuum in the synaptic input connectivity rather than distinct classes of interneurons in terms of their inputs.

### 5.2. Relation to the classical neurophysiological literature

In our data, one could see traces hinting at a classification scheme, based on input connectivity. For instance, the concept of separated Ia interneurons and Ib interneurons could have arisen as a consequence of the fact that we found convergent connections of Ia and Ib afferents to be reduced compared to chance (Figure 8). We found extensive convergence between cutaneous and proprioceptive afferents. The organization of cutaneous afferent inputs is considered to be a major remaining gap in the understanding of spinal motor systems (Loeb and Tsianos, 2015). While it has been noted that spinal interneurons can receive cutaneous inputs in addition to proprioceptive inputs, the cutaneous inputs have typically been grouped under the flexor reflex afferent (FRA) concept (Jankowska, 1992; McCrea, 1992). Hence, their contribution to the spinal cord connectivity has not been explored in great detail except for the topographical organization of cutaneous afferent inputs to the spinal cord (Levinsson et al., 2002) and the input-output relationship between skin input and controlled movement (Petersson et al., 2003). Our study could not show how input from different cutaneous afferents is organized, because we did not explore A-delta and C-fiber inputs and because we did not explore laminae I-II where these inputs dominate (Todd, 2010; Abraira et al., 2017; Häring et al., 2018).

### 5.3. Relation to classification schemes proposed based on gene expression patterns

More recently, a classification scheme for spinal interneurons has been proposed based on their gene expression associated with the initiation of spinal interneuron differentiation in embryonic development (Alstermark and Isa, 2012; Zampieri et al., 2014; Azim et al., 2014; Gabitto et al., 2016; Zholudeva et al., 2021). These genetic classes have been associated with specific functions and involvement in specific tasks. Additionally, matches between the classical neurophysiological input classification scheme with the genetic classification scheme have been suggested to exist (Brownstone and Bui, 2010; Bikoff, 2019). However, what would seem as a significant caveat to the notion of a genetically predefined wiring of the spinal interneuron circuitry, is that the equivalent of the spinal cord developed 420 million years ago, when the phylogenetic line leading to mammalians diverged from the elasmobranchs (Grillner, 2018; Jung et al., 2018). It was noted that ”*the skate spinal cord has the same interneuronal building blocks available to form the locomotor network as seen in mammals*” (Grillner, 2018), which encompasses animals of a great somatic anatomical diversity and variety of non-overlapping movement patterns. Furthermore, the spinal cord circuitry is able to adapt to such dramatic biomechanical changes as a reconfiguration of the arrangement of muscles (Loeb, 1999). Such observations indicate that if there is a genetic pre-configuration of the spinal cord circuitry, it must be possible to overwrite even during the adult life span. Further evidence is that the distribution of the primary afferent fibers is correlated with the usage of the corresponding muscles (Fritz et al., 1989).

Transcriptomics data shows that spinal interneurons in laminae V-VII, a subset of the laminae which we recorded from, do not cluster well based on their transcriptomic profile (Russ et al., 2021). This is in line with our results and another piece of evidence that learning could be an important factor in this part of the spinal cord. However, it is not clear how learning in this region of the spinal cord could be implemented at the molecular level. Transcriptomics data also shows an uneven distribution of gene expression of a subset of molecules commonly associated with learning (Russ et al., 2021). It is unclear how this gene expression varies over development, when the networks that we studied were formed. Neither is it clear how gene expression could determine patterns of convergent proprioceptive and cutaneous input connectivity. Even in the dorsal horn, developmental afferent wiring is activity dependent (Granmo et al., 2008), which indicates that even early structuring of the spinal cord circuitry is to a large extent influenced by learning and activity.

### 5.4. Implications for the functional interpretation of the spinal cord circuitry

The relatively unique input connectivity of each spinal interneuron strongly suggests that these inputs were a consequence of learning. Because of the diversity, innate, or genetically defined, patterns of input connectivity is an unlikely explanation of our results, though a combination of innate patterns with extensive modification through learning can of course not be ruled out by the present set of experiments.

An important consequence of learning is that the circuitry becomes more adaptable to the biomechanical properties of the body. A previous spinal cord model was constructed using the limited number of spinal neuron classes defined in the literature. Already this type of spinal cord system provides the possibility to generate a great diversity of temporally coherent movements using simple descending control (Raphael et al., 2010). To rebuild such a model without neuron classes and instead with a focus on a diversity of input connections would be expected to perform at least as good in this respect, because all neurons that would be representative for a class would still be present in addition to a wide diversity of other input connectivities. Such diversity of connectivities could enrich the repertoire of possible muscle activation patterns and could help resolve long standing controversies around synergy control (Tresch and Jarc, 2009). Whereas the dominant solutions in the spinal cord circuitry to descending motor commands would equal our most commonly observed movement patterns or synergies (Santello et al., 2013), a great variety of spinal interneurons would help preventing the system from getting stuck in only those movement patterns and provide the motor control systems of the central nervous system with a major potential flexibility to generate movements that are still coherent and organized.

## 6. Limitations of the Study

The study was limited by the stimulation sources. The DR nerve supplies a high number of muscles, but does not contain antagonists. We could not be certain that the inputs from Ia and Ib afferents and excitatory versus inhibitory synaptic inputs, were from the same muscle or not, and interactions between antagonists and agonists could also not be studied. In the analysis, there could be confusion between a small number of high threshold Ia and low threshold Ib inputs. The stimulation of the skin was limited to the site of maximum sensitivity. Other skin sites could have provided synaptic inputs of another type of input connectivity. Therefore our data represents an underestimate of the complexity of the input connectivity to the spinal interneurons. The analysis of the spinal network structure, especially the analysis of recurrent excitatory connections, relies on the input connections being limited to the input connections that we found.

## 7. Acknowledgments

This research received funding from the German Aerospace Center under the research and development contracts D/572/67246870, D/572/67270080, D/572/67302614 the European Union’s Horizon 2020 Framework Programme for Research and Innovation under the Specific Grant Agreements No. 720270 (Human Brain Project SGA1), 785907 (Human Brain Project SGA2), 945539 (Human Brain Project SGA3), the Hand Embodied (THE) (an Integrated Project funded by the EU under FP7, project no. 248587) and the Swedish Research Council.

## 8. Author Contribution

Conceptualization, H.J., F.B., A.A.S.; Methodology, H.J., F.B., A.A.S., M.K., P.S.; Software – Network Structure, M.K.; Software – Association Rules, M.K.; Formal Analysis, H.J., M.K., P.S.; Investigation, H.J., F.B.; Writing – Original Draft, H.J., M.K.; Writing – Review & Editing, H.J., M.K., P.S., A.A.S.; Supervision, H.J., A.A.S.; Funding Acquisition, A.A.S., F.R., H.J., A.K.;

## 11. Star Methods

### 11.1. KEY RESOURCE TABLE

| REAGENT or RESOURCE | SOURCE | IDENTIFIER |
| --- | --- | --- |
| Deposited data |  |  |
| Recording data | This study | https://doi.org/10.5281/zenodo.6257949 |
| Experimental models: Organisms/strains |  |  |
| Cats | N/A | N/A |
| Software and algorithms |  |  |
| Custom Software | This study | https://doi.org/10.5281/zenodo.6257949 |
| Spike 2 | CED | https://ced.co.uk/ |
| MATLAB | MathWork | https://www.mathworks.com/ |
| Python | Python | https://www.python.org/ |
| R | R Project | https://www.r-project.org/ |

### 11.2. RESOURCE AVAILABILITY

#### Lead contact

Further information and requests for resources should be directed to and will be fulfilled by the lead contact, Matthias Kohler.

#### Materials availability

This study did not generate new unique reagents.

#### Data and code availability

- Clustering algorithms are implemented in R using the `cluster` (Maechler et al., 2019) and the `sigclust` packages. Swap randomization, association rule and graph structure analysis are implemented in custom C++ software.
- The data and code is available at https://doi.org/10.5281/zenodo.6257949.

### 11.3. EXPERIMENTAL MODEL AND SUBJECT DETAILS

The procedures of all experiments were approved in advance by the local Swedish Animal Research Ethics Committee (permits M32-09 and M05-12). Eighteen adult cats of both sexes (> 6 months) were prepared, similar as reported previously (Jörntell and Ekerot, 2002, 2006; Spanne et al., 2014), as follows. After an initial anesthesia with propofol (Diprivan™Zeneca Ltd, Macclesfield Cheshire, UK), the animals were decerebrated at the intercollicular level and the anesthesia was discontinued. The animals were artificially ventilated and the end-expiratory CO_2_, blood pressure and rectal temperature were continuously monitored and maintained within physiological limits. Mounting in a stereotaxic frame, drainage of cerebrospinal fluid, pneumothorax and clamping the spinal processes of a few cervical and lumbar vertebral bodies served to increase the mechanical stability of the preparation. To verify that the animals were decerebrated, we made EEG recordings using a silver ball electrode placed on the surface of the superior parietal cortex. Our EEG recordings were characterized by a background of periodic 1 kHz to 4 kHz oscillatory activity, periodically interrupted by large-amplitude 7 kHz to 14 kHz spindle oscillations lasting for 0.5 s or more. These forms of EEG activities are normally associated with deep stages of sleep (Niedermayer and Lopes Da Silva, 1993). The pattern of EEG activity and the blood pressure remained stable, also on noxious stimulation, throughout experiments (see also (Jörntell and Ekerot, 2006)). Laminectomies were performed from spinal segments C3 to T1. Clamps were put on vertebra T1 and around vertebrae L1-L3. The dura was cut and a local, stabilizing coil of Teflon-coated silver wire was put on the surface of the recording area. We also made small openings in the pia mater in order to facilitate penetration of the patch clamp pipettes. The recording region was covered with agarose attached to the bone of the vertebrae in order to dampen tissue movements. See Figure 1 A for an illustration of the general experimental setup.

### 11.4. METHOD DETAILS

#### 11.4.1. Stimulations

In order to study synaptic integration in the recorded neurons, we used a set of input sources which were electrically stimulated to evoke synaptic inputs to the recorded neurons. These consisted of the deep radial (DR) nerve, for the activation of muscle afferent inputs of type Ia and of type Ib (corresponding to muscle spindle afferents and Golgi tendon organ afferents, respectively), and the skin area that by manual identification could be identified as the most sensitive for that specific neuron. The DR nerve does not contain any skin afferents but supplies the *Extensor carpi radialis brevis, Supinator, Posterior interosseous nerve* (a continuation of the deep branch after the *supinator*), *Extensor digitorum, Extensor digiti minimi, Extensor carpi ulnaris, Abductor pollicis longus, Extensor pollicis brevis, Extensor pollicis longus* and the *Extensor indicis.* A bipolar stimulation electrode was mounted around the DR nerve, which was dissected to gain access to wrap the stimulation electrode around it but otherwise leaving the DR nerve intact. The nerve and the stimulation electrodes were embedded in cotton drenched in paraffin oil to prevent spread of stimulation current into the surrounding tissue.

In order to activate A-*β* tactile afferent input from the identified skin receptive field of the recorded neuron, we inserted a pair of bipolar skin stimulation needle electrodes (separated by about 5 mm to 7 mm) into the epidermis and used single pulse electrical bipolar stimulation of an intensity between two and ten times the stimulation threshold, as explored before in a similar preparation (Bengtsson et al., 2013). This stimulation intensity corresponded to a stimulation current typically of 0.5 mA but at most 2 mA at a 0. 14 ms pulse width. At the highest of these stimulation intensities an activation of A-*δ* afferents, which typically have a conduction velocity between 1/10 and 1/5 of the A-*β* tactile afferents (Harper and Lawson, 1985; Pogatzki et al., 2002), cannot be guaranteed to be excluded. However, because their conduction velocity is more than three times slower than that of the A-*β* tactile afferents, even any possible monosynaptic response from A-*δ* afferents would have been slower than the longest latency times for the slowest, quadsynaptic inputs we observed. We did not observe any monosynaptic responses compatible with A-*δ* inputs.

#### 11.4.2. Obtaining recordings

Patch clamp pipette electrodes were pulled to impedances of 4 MΩ to 14 MΩ (Jörntell and Ekerot, 2006; Spanne et al., 2014). The pipettes were back-filled with an electrolyte solution containing potassium gluconate (135 mM), HEPES (10mM), KCl (6.0mM), Mg-ATP (2mM), EGTA (10mM). The solution was titrated to 7.35–7.40 using 1 M KOH. The recording electrodes were inserted vertically through the dorsal surface of the exposed spinal cord and spinal neurons were recorded from the lower cervical segments (C6-C8) at depths of 1.3 mm to 3.9 mm, corresponding to laminae III-VIII (Figure 1 B) (Rexed, 1952). Since all but two of the recordings were made above the depth of the alpha-motoneurons they were from putative spinal interneurons. The recording signal was amplified using the HEKA EPC 800 (HEKA, Reutlingen, Germany) and converted to a digital signal with the analog-to-digital converter Power 1401 mkII from Cambridge Electronic Design (CED, Cambridge, UK). The neural recordings were continuous and sampled at 100 kHz with the software Spike 2 from CED. The criteria used for inclusion or termination of an intracellular recording were a stable membrane potential of less than −50 mV and stable maximal peak amplitudes of spontaneous EPSPs (> 2mV, varying between neurons). In order to facilitate the analysis of evoked excitatory postsynaptic potentials (EPSPs) and inhibitory postsynaptic potentials (IPSPs), a mild (10-30 pA) hyperpolarizing bias current was injected through the recording electrode to suppress spiking activity. Before and after the neuron was recorded intracellularly, we also recorded extracellularly to allow a field potential analysis of the inputs evoked by the sensory stimulations.

We recorded from 114 spinal interneurons, with 68 spinal interneuron recordings in the whole cell mode, where the recording quality allowed the identification of the amplitude of individual synaptic responses, and thus permitted a synaptic weight analysis (Figure 2). The remainder of the recordings (*n* = 46) were not fully under whole cell control but still partially intracellular and were hence used as additional material in the analysis of the input connectivity of the peripheral inputs to the spinal interneurons.

#### 11.4.3. Analysis of PSPs

To identify a threshold stimulation intensity *T* for each response component, each input was activated at a number of different intensities, with a high number of repetitions for each intensity. All responses were analyzed using a custom-made software, which allowed a template-based identification of EPSPs and IPSPs (Figures 1 D and 2 A). These templates were created from spontaneous unitary EPSPs or spontaneous IPSPs, which were also congruent with unitary PSPs evoked at low-threshold stimulation of peripheral afferents. Evoked responses were analyzed using fitting of the template PSP shapes through scaling. The analysis aimed to define the input connectivity to each neuron. This means whether the input was mono-, di- or trisynaptic, in addition to whether it was excitatory or inhibitory.

Two types of field potentials were used to determine PSP latency. First, the afferent fiber **volley**, which is the field potential evoked by action potentials from the afferents entering the spinal cord. Second, the synaptic local field potential (**LFP**) which is caused by synaptic currents evoked by the afferent fiber volley.

Also the respective minimal threshold intensities for evoking a response were identified in this way. Response latency times were analyzed as the relative time to these field potential latency times, which made the identification of monosynaptic responses straight forward (Figure 2 D,F) (Quevedo et al., 2000; McCrea et al., 1995). In addition, monosynaptic responses could be separated from non-monosynaptic responses by their lower variability in response latency time and peak amplitude. Di-synaptic responses could be identified by their relatively minor difference of the response latency time compared to the monosynaptic response. Trisynaptic responses were separable from di-synaptic responses again in part on the differences in response latency times and relative latency to a di-synaptic response. The analysis of indirect PSP responses that were preceded by earlier PSP response components was supported by the identification of the template shapes of the EPSPs and IPSPs of each neuron. In particular, indications of rapid changes in the membrane potential that were incompatible with the expected decay phase of the template PSPs, and which was followed by the apparent addition of a new PSP component, based on template-shape identification, was taken as an indication of an another evoked PSP with a longer synaptic linkage.

#### 11.4.4. QUANTIFICATION AND STATISTICAL ANALYSIS

##### Classes of Spinal Interneurons

We used statistical tests and machine learning algorithms to determine if the spinal interneurons fell into well separated classes and to find patterns indicative of any formative process that could explain the input connectivity to the neurons. These tests did not assume a priori that a subdivision or any specific pattern must exist, hence allowing an unbiased evaluation of the input connectivity to the spinal interneurons.

In order to determine if an input-based subdivision of the neurons existed we used two conceptually similar but separate tests. The first method, SigClust(Liu et al., 2008) tests the hypothesis that the data comes from a single Gaussian distribution (Figure 5 B). To achieve this, the clusterability of the actual dataset is compared to that of data generated by sampling (*n* = 10000) from a Gaussian distribution. The Gaussian distribution was constructed by fitting it to the data, so that the mean and the variance were identical to that of the recorded data. The cost of a clustering, also called clusterability, is defined as the sum of the distances, of each datapoint to the mean of its assigned cluster. The cluster index is the fraction between the cost of clustering the data into two cluster and into one cluster. The cluster indices of the generated data are computed, giving a distribution of cluster indices. If the clustering index of the original dataset falls into the 0.05 quantil of the distribution, then the hypothesis can be rejected and the actual dataset is determined to be clusterable into at least two clusters. This method is suitable for high dimensional, low sample size data and has been applied previously in the context of biological data analysis (Liu et al., 2008; Verhaak et al., 2010; Burstein et al., 2015; Prat et al., 2010).

The second method, GapStatistics(Tibshirani et al., 2001), is conceptually similar to the SigClust method. It compares the actual data to a random distribution (Figure 5 D). Additionally, the GapStatistics determines the correct number of clusters. A possible outcome is that the number of clusters is estimated to be only one cluster or that there is as many clusters as there are datapoints. In both these cases the dataset is considered unclusterable. The GapStatistics measures how far the clustering cost of the dataset differs from the clustering cost of the reference random distribution (a uniform distribution fitted to the actual dataset) for any given number of clusters. To achieve this, random samples (*n* = 2000) are drawn from the reference distribution, and their average clustering cost is computed. Then the original data is clustered and the clustering cost is computed. The gap is the difference between the logarithmic value of the clustering cost of the actual dataset and the logarithmic value of the average clustering cost of the random samples. The number of clusters is estimated as the number which maximizes the gap. This method has also been applied to analyze biological data (Tothill et al., 2008; Grün et al., 2015; Viventi et al., 2011).

##### Associations between neuronal inputs

In order to find patterns of input connectivity we used Association Rule Mining (Agrawal et al., 1993). This method finds associations between binary properties in a given dataset. In the present context a binary property is the presence or absence of a specific input connectivity to a spinal interneuron. A potential association of two input connectivities is encoded as a so called association rule *P*_1_ ⇒ *P*_2_, where *P*_1_ and *P*_2_ are two connectivities from afferents to neurons. Such a rule is to be read as that if a neuron has connectivity *P*_1_ then it also has connectivity *P*_2_. If a neuron has connectivity *P*_1_ then the rule applies to the neuron. If a neuron has connectivity *P*_1_ and connectivity *P*_2_ then the rules applies to the neuron and holds true. If the neuron has connectivity *P*_1_ but does not have connectivity *P*_2_, then the rule applies to the neuron, but does not hold true.

For example, connectivity *P*_1_ is a monosynaptic Ia afferent input, and *P*_2_ an inhibitory disynaptic Ib afferent input. Consider four different imaginary neurons and the rule *P*_1_ ⇒ *P*_2_. Neuron one has a monosynaptic Ia afferent input and an inhibitory disynaptic Ib afferent input. Neuron two does not have Ia input but does have Ib inhibitory input. Neuron three has the Ia input but not the Ib input. Neuron four has neither input. The rule applies to neuron one and three, but does no apply to neuron two and four. The rule holds true for neuron one and does not hold true for neuron three.

For a set of neurons *S* the support of the rule *P*_1_ ⇒ *P*_2_ is

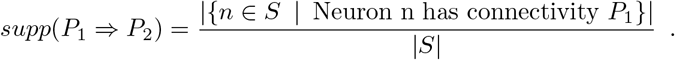

The support is the fraction of neurons where the rule applies.

The confidence of a rule

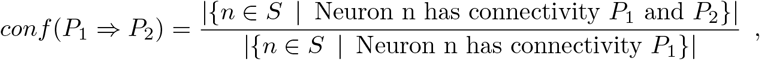

is the fraction between the number of neurons where the rule applies and where the rule holds true.

##### Graph structure analysis

We also designed a data analysis method to gain insight into the spinal interneuron network structure. Each neuron gives a glimpse of a fraction of the network connectivity. Our method works by searching for neurons whose input connectivity allows for possible connections between them. Each neuron is characterized by its input connectivity which is a subset of all possible input connections *A* = {Ia, Ib, Skin} × {-4,…, −1,1,… 4}. Here the inputs a neuron receives are encoded as a tuple consisting of the afferent where the input comes from and the number of synapses over which the input is transmitted. Inputs over more than four synapses are ignored. Inhibition is encoded as a negative number.

If a neuron *N*′ ⊆ *A* gives excitation via *n* consecutive synaptic linkages to a neuron *N*, then neuron *N* receives the following set of excitatory inputs

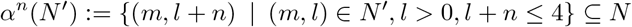

If a neuron *N*′ ⊆ *A* gives inhibition via n synapses to a neuron *N*, then neuron *N* receives the following set of inhibitory inputs

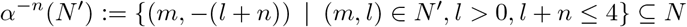

In both cases neuron *N*′ receives all excitatory input connections that *N* receives over n more synapses and in the case that *N* inhibits *N*′, *N*′ receives the input connection as inhibition.

In a network where two neurons *N*_1_, *N*_2_ ⊆ *A* are connected to each other, the following facts hold.

- If *N*_1_ and *N*_2_ are connected to each other with excitatory synapses, then *α*^+1^(*N*_1_) ⊆ *N*_2_ and *α*^+1^(*N*_2_) ⊆ *N*_1_.
- If *N*_1_ is connected to *N*_2_ with an excitatory synapse and *N*_2_ is connected to *N*_1_ with an inhibitory synapse then *α*^-1^(*N*_1_) ⊆ *N*_2_ and *α*^+1^(*N*_2_) ⊆ *N*_1_.

These facts allow us to count the number of neurons in the dataset which could be part of such a two neuron circuit, by counting the number of recorded neurons where another recorded neuron exists, so that one of the facts hold.

Each neuron receives its input either monosynaptically from the inputs or non-monosynaptically via other neurons. These neurons that transmit this input must at least receive the input they are transmitting but with one shorter synaptic linkage. These neurons, which are the minimal requirement for a predecessor neuron, to a neuron *N* are called the virtual neurons

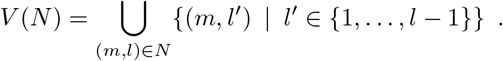

For a neuron *N* to receive its input completely over a feed forward network in contrast to transmission via network containing loops, for every virtual neuron *N*^*V*^ ∈ *V*(*N*) there must be a matching neuron *N*′ such that the following two conditions hold. The neuron *N*′ receives the input *N^V^* receives, *N^V^* ⊆ *N*′. If *N^V^* transmits its input over *n* synapses to *N* then *N* must contain all inputs that *N*′ receives transmitted over n synapses *α*″(*N*′) if the transmitted input at *N* is excitatory or *α*^-1^(*N*′) if the transmitted action at *N* is inhibitory. This kind of analysis relies on the assumption that only the patterns of convergence that we found exist. It is unlikely that the complete set of connectivity within the spinal cord circuitry was mapped in this study.

##### Swap Randomization

In order to determine if the associations and graph structures quantified by the measures described in the previous sections are by chance, we compared the measures to datasets generated by swap randomization. The swap randomized datasets are created by virtually swapping inputs between neurons under the following constraints, as previously described(Gionis et al., 2007). Each neuron receives the same number of inputs as in the original dataset. Each input contacts the same number of neurons as in the original dataset. If the measure obtained from the recorded dataset is smaller than the average of the measure on the swap randomized datasets, we call the measure reduced. In the opposite case we call it enhanced. Let *M* be the value of a measure computed on the recorded dataset. Let *a* and *b* be the number of swap randomized datasets where the measure is smaller than *M* respectively larger than *M* The significance of the comparison to the swap randomized datasets is computed as the empirical *p*-value

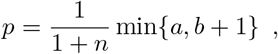

where *n* = 10^7^ is the number of swap randomized datasets.

